# Expanded Genomic Sampling of the Desulfobulbales Reveals Distribution and Evolution of Sulfur Metabolisms

**DOI:** 10.1101/2021.02.08.430318

**Authors:** Lewis M. Ward, Emma Bertran, David T. Johnston

## Abstract

The reconstruction of modern and paleo-sulfur cycling relies on understanding the long-term relative contribution of its main actors; these include microbial sulfate reduction (MSR) and microbial sulfur disproportionation (MSD). However, a unifying theory is lacking for how MSR and MSD, with the same enzyme machinery and intimately linked evolutionary histories, perform two drastically different metabolisms. Here, we aim at shedding some light on the distribution, diversity, and evolutionary histories of MSR and MSD, with a focus on the Desulfobulbales as a test case. The Desulfobulbales is a diverse and widespread order of bacteria in the Desulfobacterota (formerly Deltaproteobacteria) phylum primarily composed of sulfate reducing bacteria. Recent culture- and sequence-based approaches have revealed an expanded diversity of organisms and metabolisms within this clade, including the presence of obligate and facultative sulfur disproportionators. Here, we present draft genomes of previously unsequenced species of Desulfobulbales, substantially expanding the available genomic diversity of this clade. We leverage this expanded genomic sampling to perform phylogenetic analyses, revealing an evolutionary history defined by vertical inheritance of sulfur metabolism genes with numerous convergent instances of transition from sulfate reduction to sulfur disproportionation.

## Introduction

Microbial sulfur metabolisms drive the biogeochemical sulfur cycle over geologic timescales and couple it to the carbon, oxygen, and iron cycles (Johnston et al., 2005; Fike et al., 2015). Though the contribution of these metabolisms to net global carbon fixation rates is low relative to that of photosynthesis (e.g., Ward and Shih 2019), carbon and sulfur fluxes through dissimilatory sulfur metabolisms are large and provide a significant control on net oxidation-reduction (redox) balance, in turn driving changes in Earth surface conditions (Berner and Raiswell, 1984; Berner and Canfield, 1989; Canfield, 2001; Fike et al., 2015). The main microbial metabolisms that drive the sulfur cycle are microbial sulfate reduction (MSR), sulfide oxidation (SO), and sulfur disproportionation (MSD) (Johnston et al., 2005; Fike et al., 2015). MSR couples the oxidation of simple organic molecules – including H_2_ for some organisms – to the reduction of sulfate, thiosulfate, and in some cases sulfite (Rosenberg et al., 2014, and references within). This reductive sulfur reaction promotes the burial of sedimentary pyrite and the remineralization of organic matter, which are major controls on Earth’s surface redox conditions (Canfield, 2001; Fike et al., 2015; Canfield and Teske, 1996; Jorgensen, 1982). MSD, heavily involved in the oxidative sulfur cycle (Canfield, 2001), is a chemolithotrophic process by which sulfur species of intermediate valence – thiosulfate, sulfite, and/or elemental sulfur – act as both electron acceptor and donor, producing sulfate and sulfide as final products (Canfield and Thamdrup, 1994; Canfield et al., 1998; Finster et al., 1998; Finster et al., 2013; Frederiksen and Finster, 2003; Habicht and Canfield, 1998; Thamdrup et al., 1993; Finster, 2008 for a review on this metabolic pathway).

It has long been understood that MSR and MSD share core reactions and enzymatic machineries (Frederiksen and Finster, 2003), yet yield drastically different net pathways - a mechanistic argument for this conundrum is lacking. MSR is a respiratory pathway while, MSD is fermentative and the energetic yields for each differ significantly. Sulfate reduction is a vastly more thermodynamically favorable metabolism than sulfur disproportionation under standard conditions (Finster, 2008; Wing and Halevy, 2014). As an example, in the absence of a sulfide sink, elemental sulfur disproportionation is effectively an endergonic reaction (Finster et al., 1998; Finster, 2008). It was then surprising when early pure culture experiments, as well as full genome sequencing and enzyme extract studies, revealed sulfate reduction and sulfur disproportionation share the same sulfur metabolism enzymes – sulfate adenylyltransferase, adenylylsulfate reductase (subunits A and B), dissimilatory sulfite reductase (subunits A, B, and C), and the sulfite reduction-associated DsrMKJOP complex – (Frederiksen and Finster, 2003; Finster el al., 2013). These are also dramatically different from those enzymes driving sulfide oxidation. It would thus be expected for sulfur disproportionating microbes to be capable of using sulfate as an electron acceptor in the presence of organic matter, and for sulfate reducers to disproportionate sulfur species of intermediate valence when conditions permitted. This expectation on the metabolic plasticity of sulfate reducers and sulfur disproportionators is not met and most sulfur disproportionators are incapable of MSR (Finster et al., 1998). To date, only two exceptions to this phenomenon have been reported: *Desulfocapsa thiozymogenes* (Junghare and Schink, 2015; Rosenberg et al., 2014; Canfield *et al*., 1998; Janssen *et al*., 1996) and *Desulfobulbus propionicus* (Widdel, 1980; Sorokin et al., 2012; Widdel and Pfennig,1982; Janssen et al., 1996; Böttcher et al., 2005; El Houari et al., 2017; Kramer and Cypionka 1989; Lovley and Phillips 1994; Fuseler and Cypionka 1995).

Understanding the similarities and differences between MSR and MSD carries geological importance. The antiquity of sulfate reduction has been dated back to the Archean using the MSR sulfur isotopic signature (Shen et al., 2001; Bontognali et al., 2012). On the other hand, the antiquity of sulfur disproportionation is harder to pinpoint. Chemical and isotopic signatures suggest the rise to ecological significance of MSD to be as late as the Mesoproterozoic (Johnston et al., 2005) or as early as the Archean (Phillippot et al., 2007), and molecular clock work to refine the timing of emergence of MSD is lacking. For thermodynamic reasons, the ecological niche occupied by MSD includes partially oxidizing conditions and a sulfide sink (Finster, 2008), conditions that are met only post the Great Oxidation Event. This would then suggest that MSD rose to ecological significance later than MSR (Canfield and Teske, 1996). All in all, the shared nature of the MSR and MSD metabolic pathways coupled with full genome sequencing efforts have led to the inference that MSR and MSD are old and share a complex evolutionary history that is difficult to untangle.

The knowledge gap here resides with our understanding of MSD, as MSR has been far more thoroughly studied (Fry et al., 1986; LeGall and Fauque 1988; Canfield, 2001; Shen et al. 2001; Habicht and Canfield, 1998; Habicht et al., 2002; Zane et al, 2010; Pereira et al., 2011; Keller and Wall, 2011; Leavitt et al;, 2013; Parey et al., 2013; Wing and Halevy, 2014; Fike et al., 2015; Bradley et al., 2016; Bertran et al., 2018). That is, the true diversity and ecological distribution of sulfur disproportionation is still unknown owing to the lack of a unique enzymatic and genetic marker. Recently, increased efforts, technological advancements and sampling in metagenomics have expanded the ecological distribution and significance of MSR in modern sediments (Anantharaman et al., 2018; Vigneron et al., 2018). However, and as noted above, efforts for MSD are lagging due largely to the absence of established marker genes to distinguish the capacity of MSD from MSR based on genome data alone (Anantharaman et al., 2018). The true diversity and ecological distribution of S metabolisms is still poorly understood despite the central role of these pathways in modern and past biogeochemical cycling and potential role in neo-environments as sulfidic environments spread with changing climate.

An ideal case study for investigating the evolutionary relationship between MSR and MSD exists in the bacterial order Desulfobulbales. The Desulfobulbales are members of the Desulfobacterota (formerly Deltaproteobacteria) phylum (Fig. 2) and include diverse and environmentally widespread members that play a central role in sulfur biogeochemical cycling in both modern and paleo-sediments (Fike et al., 2015) (Fig. 1). The Desulfobulbales were first described in 1980 when Widdel and colleagues described Desulfobulbus, the type genus of the order (Widdel et al., 1980; Kuever et al., 2005), but now consist of at least three family-level clades spanning at least ten genera (Fig. 3). Members of the Desulfobulbales order have been described by a wide a range of morphological and chemotaxonomic properties (Rosenberg et al., 2014), and while they have been isolated from various sources – freshwater, marine environments, brackish water, and haloalkaline environments – most are mesophilic bacteria and all isolates are strictly anaerobic (Kuever, 2014). The Desulfobulbales order also includes the recently discovered filamentous “cable bacteria” (Kieldsen et al., 2019), which have been shown to link redox processes across sediment layers separated by distances over 1 cm via long-distance electron transport (Müller et al., 2016) and may even be capable of sulfur disproportionation under some conditions (Müller et al., 2020). However, cable bacteria have so far resisted isolation in pure culture, preventing detailed physiological characterization (Pfeffer et al., 2012; Schauer et al., 2014; Bjerg et al., 2016; Kjeldsen et al., 2019). As a result, cable bacteria will not be included in our analysis.

**Figure 1:**
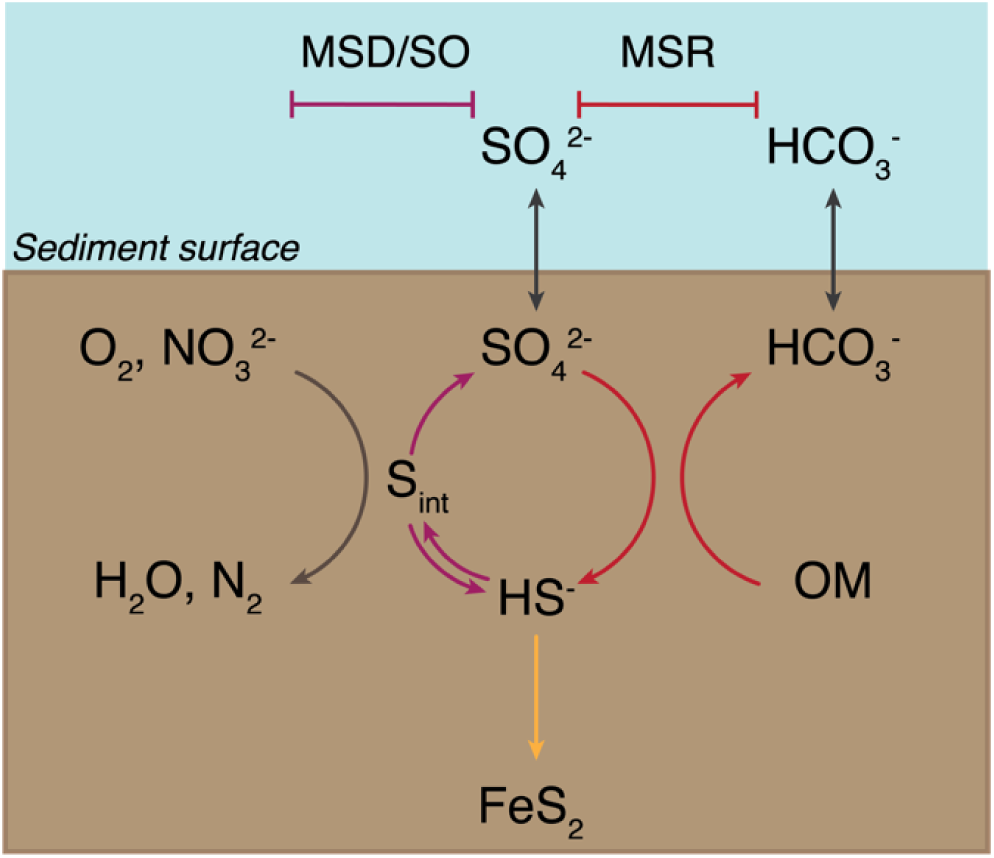
Sedimentary biogeochemical sulfur cycle. The sediment-water interface is indicated at the “Sediment surface”. First, seawater sulfate (SO_4_^2-^) diffuses into the sediments and enters the reductive sulfur cycle promoted by microbial sulfate reduction (MSR) (indicated with red arrows), which couples the reduction of sulfate to sulfide (HS^-^) to the oxidation of organic matter (OM). A fraction of the produced sulfide precipitates with iron to form pyrite (FeS_2_) and is ultimately buried (yellow arrow). A portion of biogenic sulfide is oxidized – either biotically through sulfide oxidation (SO, shown with a blue arrow) or abiotically – using common oxidants – oxygen (O_2_) or nitrate (NO_3_^2-^) – to yield intermediate sulfur species (S_int_). These are then disproportionated via microbial sulfur disproportionation (MSD) to release sulfate and sulfide (depicted with a purple arrow).

**Figure 2:** **(A)** Tree of Life built with concatenated ribosomal proteins following Hug *et al*. (2016) collapsed at the phylum level as classified by GTDB-Tk showing the relationship of Desulfobacterota relative to Proteobacteria and other major bacterial groups. **(B)** Concatenated ribosomal protein phylogeny of the Desulfobacterota binned at the family (Desulfobulbales) or class (all other lineages) levels, labeled with taxonomic assignments from GTDB-Tk, showing the placement of and relationships within the Desulfobulbales. Nodes are labeled with TBE support values.

**Figure 3:** Phylogenetic tree showing the genomes of isolated and well-characterized members of the Desulfobulbales, including the families Desulfobulbaceae, Desulfocapsaceae, and Desulfurivibrionaceae. Nodes are labeled with TBE support values. Species names are highlighted with colors corresponding to the taxonomic family to which they are assigned. On the right, the characterized capacity for performing sulfur metabolisms is indicated.

**Figure 4:**
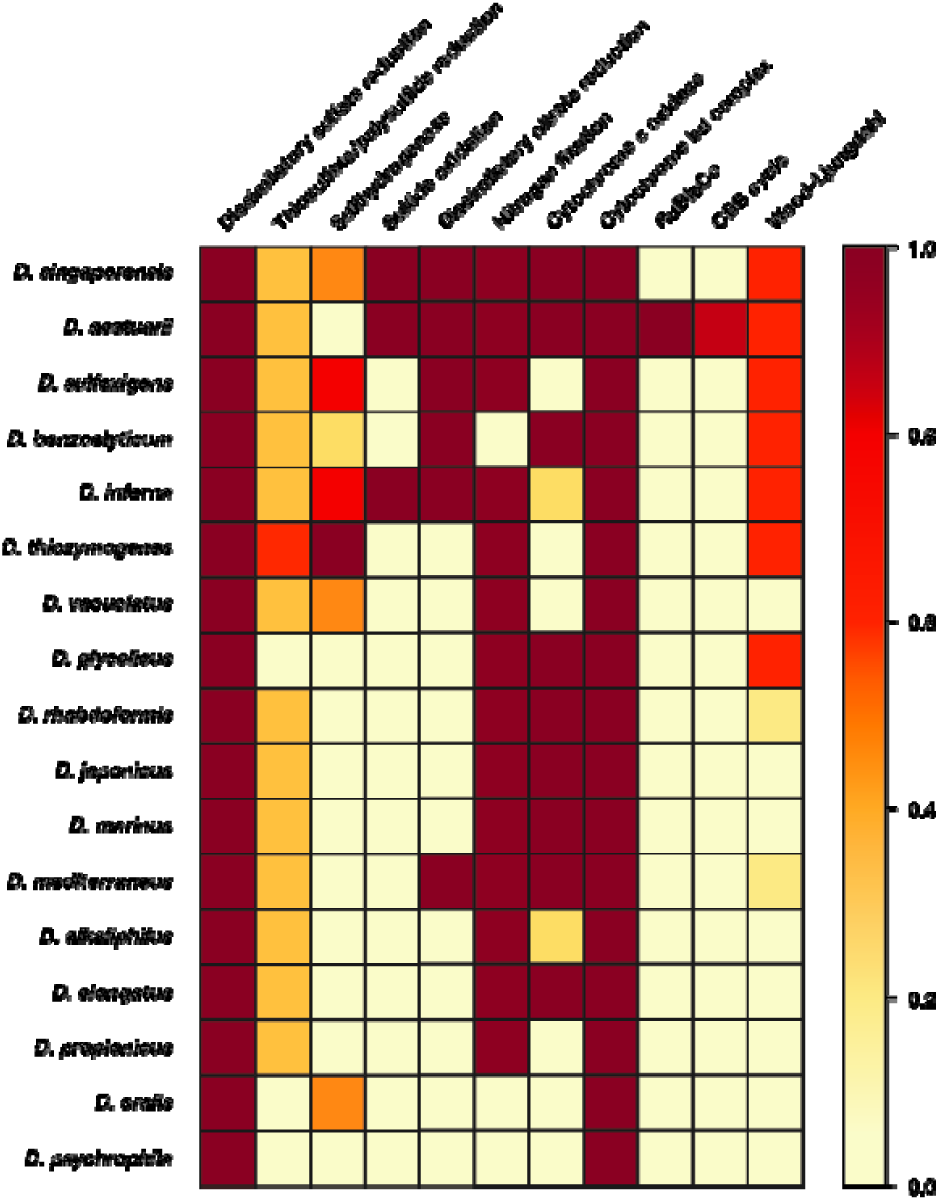
Heatmap of metabolic functions produced by the KEGG-decoder of the members of the Desulfobulbales sequenced here. The color gradient reflects the fractional abundance of genes associated with a pathway encoded by a particular genome. In other words, white implies no genes associated with a pathway of interest are found in the genome and thus that said pathway is no constituted. Conversely, dark red indicates all genes required to perform the pathway of interest are found and that said metabolism is fully constituted in the genome. Implications for the presence or absence of metabolic pathways of interest in each genome are discussed in the text.

Here, we first provide a revised genome-based taxonomy of the Desulfobulbales. We do this by presenting draft genomes of previously unsequenced isolates belonging to the Desulfobulbales, and couple this newly expanded genomic sampling with comparative genomic and phylogenetic analyses. We then examine the distribution and potential ecological diversity of MSR and MSD and provide insight into the relationships between strains previously characterized as sulphate reducers or sulfur disproportionators. Ultimately, these will provide a refined assessment of the major evolutionary transitions between the lineages of sulfate reduction and sulfur disproportionation.

## Methods

### Genome sequencing and analysis

Our analyses focused on Desulfobulbales strains that have both been well characterized in pure culture and for which high-quality genome sequences are available. We therefore omitted organisms known only from metagenome-, environmental-, or enrichment-based analyses, including the cable bacteria. Preexisting genome sequences of Desulfobulbales were downloaded from the NCBI WGS and Genbank databases. In order to thoroughly sample the genomic diversity of well-characterized Desulfobulbales isolates, we also performed genome sequencing on six species of Desulfobulbales that are available in pure culture but for which genome sequencing had not previously been performed. These included: *Desulforhopalus vacuolatus* (Isaksen and Teske 1996), *Desulfobulbus marinus* (Widdel and Pfennig 1982, Kuever et al., 2015), *Desulfoprunum benzoelyticum* (Junghare and Schink 2015), *Desulfopila inferna* (Gittel et al., 2010), *Desulfobulbus rhabdoformis* (Lien et al., 1998), and *Desulfobulbus alkaliphilus* (Sorokin et al., 2012).

Draft genomes for Desulfobulbales isolates were sequenced following methods described previously (Bertran et al., 2020a, Bertran et al., 2020b, Bertran et al., 2020c, Ward et al., 2020a, Ward et al., 2020b) and outlined briefly below. Purified genomic DNA was acquired for each strain from the DSMZ and submitted to MicrobesNG for sequencing. DNA extraction was performed with a JetFlex genomic DNA purification kit from Genomed. DNA libraries were prepared using Nextera XT library prep kits using a Hamilton Microlab Star automated liquid handling system. Sequencing was performed with an Illumina HiSeq using a 250 base pair paired-end protocol. Reads were adapter trimmed with Trimmomatic 0.30 (Bolger et al., 2014). De novo assembly was performed with SPAdes version 3.7 (Bankevich et al., 2012). Genomes were annotated and analyzed using RAST v2.0 (Aziz et al., 2008). Completeness and contamination/redundancy of genomes was estimated with CheckM v1.0.12 (Parks et al., 2015). The likelihood for presence or absence of metabolic pathways was determined using MetaPOAP v1.0 (Ward et al., 2018a). Taxonomic assignments were verified with GTDB-Tk v0.3.2 (Parks et al., 2018). Hydrogenase proteins were classified with HydDB (Søndergaard et al., 2016).

### Phylogenetic analyses

Phylogenetic analyses followed methods described elsewhere (Ward et al., 2020c, Ward and Shih 2021) and summarized below. Genomes were downloaded from the NCBI Genbank and WGS databases. The dataset used for comparative genomics analyses consisted of all complete or high quality (following current standards, Bowers et al., 2017) genomes of isolated members of the Desulfobulbales. Phylogenetic analyses incorporated all genomes of isolates as well as metagenome-assembled genomes of members of the Desulfobacterota (Deltaproteobacteria) available on the NCBI Genbank and WGS databases as of August 2019. Protein sequences used in analyses (see below) were identified locally with the *tblastn* function of BLAST+ v2.6.0 (Camacho et al., 2009), aligned with MUSCLE v3.8.31 (Edgar, 2004), and manually curated in Jalview v2.10.5 (Waterhouse, 2009). Positive BLAST hits were considered to be full length (e.g. >90% the shortest reference sequence from an isolate genome) with e-values better than 1e-20. Phylogenetic trees were calculated using RAxML v8.2.12 (Stamatakis, 2014) on the Cipres science gateway (Miller et al., 2010). Transfer bootstrap support values were calculated by BOOSTER (Lemoine et al., 2018). Trees were visualized with the Interactive Tree of Life viewer (Letunic and Bork, 2016). Taxonomic assignment was confirmed with GTDB-Tk v0.3.2 (Parks et al., 2018, Chaumeil et al., 2019). Amino Acid Identity of genomes was determined following methods from Rodriguez and Konstantinidis (2014). The shared evolutionary histories of the sulfate reduction and sulfur disproportionation lineages was inferred by inspection of the topological congruence of organismal and metabolic protein phylogenies following previously established methods (Doolittle, 1986; Ward et al., 2018b) and following the most parsimonious conclusion on vertical inheritance, divergence, and general evolutionary transitions. Protein sequence annotation was done by GhostKOALA using default settings (Kanehisa et al., 2016) and amino acid sequences translated by Prodigal (Hyatt et al., 2010). Visualization of the presence or absence of complete or partial metabolic pathways was done using KEGG-decoder (Graham et al., 2018) after manual formatting of GhostKOALA output.

## Results and discussion

### Draft genomes of Desulfobulbales isolates

In order to improve coverage of genomic diversity of Desulfobulbales, we sequenced draft genomes from isolates of six species from the Desulfobulbaceae and Desulfocapsaceae families: *Desulforhopalus vacuolatus* (Isaksen and Teske, 1996), *Desulfobulbus marinus* (Widdel and Pfennig, 1982; Kuever et al., 2015), *Desulfoprunum benzoelyticum* (Junghare and Schink, 2015), *Desulfopila inferna* (Gittel et al., 2010), *Desulfobulbus rhabdoformis* (Lien et al., 1998), and *Desulfobulbus alkaliphilus* (Sorokin et al., 2012). Genome statistics are summarized in Table 1 and presence of relevant functional genes is described below. *D. vacuolatus, D marinus, D. benzoelyticum, D. inferna, D. rhabdoformis*, and *D. alkaliphilus* all encode the full enzymatic machinery shared by dissimilatory sulfate reduction and sulfur disproportionation, that is, DsrAB, DsrMKJOP, and AprAB (Keller and Wall, 2011; Pereira et al., 2011; Parey et al., 2013). While all these strains have been reported as sulfate reducing bacteria based on pure culture experiments geared to test their metabolic capacities (Isaksen and Teske, 1996; Widdel and Pfennig, 1982; Kuever et al., 2015; Junghare and Schink, 2015; Gittel et al., 2010; Lien et al., 1998; Sorokin et al., 2012) only, *D. vacuolatus* has been reported incapable of sulfur disproportionation (Isaksen and Teske, 1996). The other strains have yet to be tested for the capacity to disproportionate intermediate valence sulfur species. Further, a correlation between the length of the AprB C-terminus and the capacity to perform sulfate reduction or disproportionation has recently been suggested, where a truncated C-terminus would be indicative of sulfur disproportionation (Bertran, 2019). *D. finferna*, encodes a full length AprB protein, whereas *D. vacuolatus, D marinus, D. benzoelyticum, D. rhabdoformis*, and *D. alkaliphilus* encode a truncated C-terminal AprB domain like other sulfur disproportionators in the Desulfobulbaceae (Bertran, 2019). However, further work is needed to confirm the validity of this truncation as a distinct genetic marker for sulfur disproportionation and there is, to date, no definite feature that distinguishes sulfate reducers from sulfur disproportionators.

**Table 1:** Genome statistics

Despite their characterization as obligate anaerobes, *D. vacuolatus* and *D. rhabdoformis* encode the capacity for O_2_ reduction via a *bd* O_2_ reductase. While *bd* O_2_ reductase enzymes are sometimes coupled to aerobic respiration (e.g., in *Nitrospira*, Palomo et al., 2018), they can be found in obligate anaerobes (e.g., Ward et al., 2015) where they are likely associated with O_2_ detoxification and oxidative stress tolerance (Forte *et al*., 2017). Additionally, *D. benzoelyticum, D. rhabdoformis* and *D. marinus* also encode A-family heme copper oxidoreductases (HCOs) for O_2_ reduction; while these enzymes are typically coupled to aerobic respiration they can also be found in obligate anaerobes (e.g. Pace et al., 2015, Hemp et al., 2015). In anaerobic organisms, such as members of the Desulfobulbales, these proteins are likely also associated with O_2_ detoxification and oxidative stress tolerance (e.g. Forte et al., 2017). Closely related A-family HCO proteins are also encoded by other Desulfobulbales such as *Desulfopila aestuarii, Desulfofustis glycolicus, Desulfobulbus japonicus, Desulfobulbus mediterraneus*, and *Desulfobulbus elongatus*. These Desulfobulbales HCOs form a closely related clade in broad HCO phylogenies sampling across diverse bacteria and archaea (Supplemental Figure 1). This suggests broad vertical inheritance of HCOs from a common ancestor of the Desulfobulbales, with perhaps a small amount intra-order HGT, potentially suggesting a long history of aerotolerance in the Desulfobulbales that stands in contrast to the O_2_ sensitivity of other orders of Desulfobacterota (e.g., Rosenberg et al., 2014). This may have served as a preadaptation to marginal redox environments in which the transition from MSR to MSD may have been favored, leading to the relatively high density of novel transitions to sulfur disproportionation in the Desulfobulbales.

Several of the Desulfobulbales genomes reported here also encode proteins involved in nitrogen redox reactions. Nitrogen fixation via a molybdenum nitrogenase is encoded by *D. vacuolatus, D. inferna*, and *D. marinus. D. rhabodoformis* encodes both a molybdenum nitrogenase as well as a vanadium alternative nitrogenase. Additionally, *D. marinus* encodes nitrite reduction to ammonium via NrfH. Despite being characterized as incapable of nitrate respiration (Junghare and Schink, 2015), *D. benzoelyticum* encodes a pathway for nitrate reduction to ammonia including nitrate reductase and cytochrome c552 nitrite reductase.

Members of the Desulfobulbales utilize various electron donors for growth, typically including simple alcohols, organic acids, and other small organic compounds which are typically incompletely oxidized (producing CO_2_ and acetate) (Kuever, 2014). Among notable exceptions to this trend in the Desulfobulbales strains discussed here, *D. benzoelyticum* is known to completely degrade benzoate to CO_2_ (Junghare and Schink, 2015). Benzoate degradation is known to be performed by a pathway consisting first of benzoate-CoA ligase and downstream enzymes including benzoyl-CoA reductase and benzoyl-CoA 2,3-epoxidase. The *D. benzoelytcium* genome recovered genes encoding for benzoate-CoA ligase but not known genes for downstream steps. Given the high completeness of the *D. benzoelyticum* genome (∼99.85 %), the probability that the complete genome encodes additional benzoate degradation genes is incredibly low (<10^−7^) as determined by MetaPOAP (Ward et al., 2018). This suggests that *D. benzoelyticum* may utilize a novel pathway for benzoate metabolism using previously uncharacterized genes, though additional genetic and biochemical study will be necessary to validate this hypothesis.

### Diversity and taxonomy of Desulfobulbales

Classically, the family Desulfobulbaceae within the Desulfobacterales order of the Deltaproteobacteria phylum has included the genera *Desulfobulbus, Desulfocapsa, Desulfofustis, Desulfopila, Desulforhopalus, Desulfotalea*, and *Desulfurivibrio* (Kuever, 2014). However, recent attempts at more systematic and normalized taxonomies based on full genome comparisons (e.g. Rinke et al., 2020, Waite et al., 2020) provide an opportunity to reassess this classification. The Deltaproteobacteria in particular have proven excellent cases for the necessity of taxonomic reappraisal as lineages assigned to this phylum have been shown to be polyphyletic, not closely related to other groups defined as Proteobacteria, and likely to represent several phylum-level groups (e.g., Hug et al., 2016, Waite et al., 2020). In recognition of these facts, the Genome Taxonomy Database (GTDB) has divided the Deltaproteobacteria into several monophyletic phyla including Desulfobacterota, which contains the bulk of classical Deltaproteobacteria such as *Desulfovibrio, Desulfobacter*, and *Desulfobulbus* (Parks et al., 2018, Waite et al., 2020). Additionally, the GTDB proposes further subdivision of lower taxonomic levels in order to remove poly- or para-phyletic groupings and normalize taxonomic ranks (Parks et al., 2018). In the case of the Desulfobulbaceae, the GTDB has reassigned these organisms to four families (Desulfobulbaceae, Desulfocapsaceae, Desulfurivibrionaceae, and BM004) within the new order Desulfobulbales of the Desulfobacterota phylum.

Our expanded phylogeny of the Desulfobulbales is broadly consistent with the revised GTDB taxonomy (Figure 2, Figure 3), recapitulating a monophyletic Desulfobulbales order within the Desulfobacterota as well as producing consistent family-level groupings within this order. The GTDB further suggests subdivision of the *Desulfobulbus* genus into at least two genera within the Desulobulbaceae family. AAI analyses (Supplemental Table 2) shows no higher than 75% similarity in any pairwise comparison of characterized Desulfobulbales strains, consistent with each strain representing at least a unique species. Genus-level cutoffs of 55-60% largely follow taxonomic boundaries assigned based on physiology and other classical metrics. While the GTDB suggests the subdivision of *Desulfobulbus* into at least two genera – that is, *Desulfobulbus sensu stricto* which includes *D. marinus, D. oralis, D. propionicus*, and a genus including *D. rhabdoformis* and *Desulfobulbus A* containing *D. japonicus* and *D. mediterraneus* – this subdivision is only somewhat supported by AAI analyses. Pairwise AAI similarity between *Desulfobulbus* strains is only < 0.55 for *Desulfobulbus oralis* when compared against *D. japonicus, D. mediterraneus*, or *D. marinus*. Pairwise comparisons between other members of *Desulfobulbus sensu stricto* and *Desulfobulbus A* largely show AAI values in the range of 0.6-0.7, consistent with a single *Desulfobulbus* genus. This, together with generally poor support for the phylogenetic placement of *D. oralis*, suggests that the relatively high divergence of *D. oralis* from other members of *Desulfobulbus* may artificially inflate the apparent taxonomic breadth of strains classified as *Desulfobulbus*. It is currently unclear whether *D. oralis* shows particularly high divergence given factors relating to adaptation to it unique niche (for *Desulfobulbales* strains) in the human mouth, because of elevated rates of mutation or horizontal gene transfer (HGT), or for other reasons. In summary, our results support the reassignment of the Desulfobulbales to the new taxonomic classification proposed by the GTDB, particularly at the family level and above. We therefore use GTDB-based clade names (e.g. Desulfobacterota, Desulfobulbales) throughout.

### Congruence of organismal and sulfur metabolic protein phylogenies in the Desulfobulbales

The distribution of sulfur metabolisms in the Desulfobulbales is scattered, with the capacity for reduction and disproportionation reactions interspersed in different groups (Figure 3). The capacity for sulfur disproportionation in particular appears to be polyphyletic. As a result, it is impossible to confidently assert a simple evolutionary history for sulfur metabolisms in the Desulfobulbales. Viable scenarios for the history of sulfur metabolisms in this clade could include, for instance, (1) an ancestor capable of both sulfate reduction and sulfur disproportionation followed by loss of either metabolism in many lineages, (2) an ancestor capable of sulfate reduction but not sulfur disproportionation, followed by convergent evolution of sulfur disproportionation, with or without loss of sulfate reduction, in many lineages and independently, or (3) the presence of sulfate reduction but not sulfur disproportionation, followed by a single evolutionary origin of sulfur disproportionation and ensuing HGT to distribute this metabolism into multiple lineages. More complicated scenarios involving multiple origins, losses, and horizontal transfers of pathways are also conceivable. Distinguishing between these scenarios is challenging, particularly given the inability to distinguish between the capacity for sulfate reduction and sulfur disproportionation via genome content alone (Anantharaman et al., 2018). The capacity for sulfate reduction and sulfur disproportionation is currently determined only thorough culture-based characterization; however, the capacity for disproportionation metabolisms is frequently not determined or reported (Figure 3). As a result, our ability to interpret the evolutionary history of sulfur metabolisms in the Desulfobulbales is limited. However, sufficient data is available to draw some conclusions about overall trends.

It is well established that sulfur disproportionation utilizes the same basic biochemical pathways as sulfate reduction, albeit with modifications to enzymes or regulation that allows some steps to run in reverse (Finster, 2008; Frederiksen and Finster, 2003; Finster et al., 1998; Finster et al., 2013). Patterns of vertical versus horizontal transfer of components in this pathway should reflect vertical versus horizontal inheritance of the metabolisms themselves. We therefore applied methods comparing organismal to functional protein phylogenies to investigate whether HGT of sulfur metabolizing proteins was responsible for the scattered distribution of sulfur disproportionation in the Desulfobulbales. If sulfur metabolism proteins (e.g. AprA, DsrA) phylogenies differ from organismal phylogenies (as determined by concatenated ribosomal proteins or other markers), this would suggest a history of horizontal gene transfer. Instead, it appears that sulfur metabolizing proteins have been vertically inherited within the Desulfobulbales, with few, if any, instances of horizontal gene transfer (Supplemental Figure 2). Rather, this supports scenarios of multiple instances of convergent evolution of sulfur disproportionation or, alternatively, the capacity for both sulfur disproportionation and sulfate reduction in the last common ancestor of the Desulfobulbales followed by many instances of loss of one pathway. The absence of intra-order HGT of sulfur metabolism pathways is further supported by the scattered but consistent localization of sulfur metabolisms genes across Desulfobulbales genomes, preventing straightforward HGT of a single operon or cluster of genes, but broadly retaining position of particular genes in the genome between members of the Desulfobulbales (e.g. colocalization of *aprAB* with the anaerobic respiratory complex *qmoABC*).

While the antiquity of sulfur disproportionation is not entirely clear, the simplest explanation for the distribution of sulfate reduction is that this metabolism was present in the last common ancestor of the Desulfobulbales and was secondarily lost in a few lineages (e.g. *Desulfocapsa sulfexigens*). This scenario is particularly compelling given the broad distribution of sulfate reduction and the relatively sparse distribution of sulfur disproportionation in the Desulfobacterota (e.g. Anantharaman et al., 2018). Whether sulfur disproportionation arose multiple times in different Desulfobulbales lineages or originated once in the stem group of this clade, it appears to represent convergent evolution with disproportionators in other lineages of Desulfobacterota and other phyla.

## Conclusion

The distribution and evolutionary history of MSR and MSD in the Desulfobacterota, and in microbes in general, is a complex palimpsest of vertical inheritance, occasional horizontal gene transfer, and extensive convergent evolution. The expanded genomic diversity of the Desulfobulbales order presented here provides additional context for investigating transitions between MSR and MSD but is unable to resolve a simple evolutionary history for this process. While it has long been apparent that sulfur disproportionation is derived from sulfate reduction, there still exists no unambiguous molecular markers to distinguish the capacity for these metabolisms from genomic data alone, nor is it clear what ecological or evolutionary processes underlie the innovation of sulfur disproportionation with or without the concurrent loss of sulfate reduction. However, the expanded genomic diversity presented here for well-characterized isolates, coupled with comparative phylogenetic approaches, can provide significant insight into the history of the Desulfobulbales. In this group, it is clear that the ancestral phenotype is of sulfate reduction, with multiple, convergent transitions to sulfur disproportionation either with or without the concurrent loss of sulfate reduction. This is in line with earlier work that supported the derivation of MSD from MSR (Canfield and Teske, 1996; Habicht and Canfield, 1998; Shen et al., 2001; Johnston et al., 2005; Philippot et al., 2007; Fike et al., 2015). By demonstrating the vertical inheritance of sulfur metabolic genes in the Desulfobulbales, we can rule out a major role for horizontal gene transfer in the distribution of MSD across the diversity of this clade. While the precise biochemical mechanisms and ecological triggers for the transition from MSR to MSD in this clade are still unknown, the propensity for the Desulfobulbales to invent and reinvent MSD may be related to a genomic background that includes pre-adaptations to marginal redox environments (e.g. presence of pathways for O_2_ detoxification) as well as alleles that allow more ready reversibility of key enzymes (e.g. the truncated AprB tail; Bertran, 2019). Further determination of markers for MSD in the Desulfobulbales and other organisms will require more thorough characterization and reporting of the capacity for disproportionation in sulfate reducing strains to reduce the burden of missing data (e.g. Fig. 3) and to better allow thorough comparative genomics to identify genetic differences between disproportionator and non disproportionator lineages.

The apparent phenotypic plasticity between MSR and MSD over relatively short evolutionary timescales (i.e. species- or genus-level variability, versus evolution over family or higher longer timescales as is typically seen in other metabolic traits like phototrophy and carbon fixation, e.g. Shih et al. 2017, Ward and Shih 2020) has significant implications for our understanding of the roles of these metabolisms in Earth history. If sulfate reducing microbes can readily and independently evolve the capacity for disproportionation, this suggests that this process may occur frequently in diverse lineages over geologic time. As a result, it is likely that sulfur disproportionating microbes have been present for as much of Earth history as there have been appropriate redox gradients in marine sediments and other environments — but, importantly, these likely have consisted of different, unrelated lineages at different times in Earth history. It is therefore reasonable to assume the activity of MSD in shaping sulfur isotopes and other sedimentary records from periods of Earth’s past, but it may not be possible to assume taxonomic affinity or other traits of the organisms responsible.

While expanded genomic sampling of the Desulfobulbales can improve our current understanding of the taxonomic and phylogenetic relationships in this clade, it is insufficient to fully untangle trends and processes in the evolutionary relationships between MSR and MSD. A major barrier to our understanding of these processes is our inability to distinguish the capacity for these metabolisms from genome content alone. Isolation and extensive physiological characterization of the sulfur metabolism capacity for Desulfobulbales strains continues to be essential; this includes the successful isolation of novel organisms in this clade (e.g. the enigmatic cable bacteria) but also the thorough testing and reporting of the sulfur disproportionation capacity for existing isolates (i.e. filling in the extensive “Not Reported” entries in Table 1 and Figure 3). Alternatively, identifying robust and consistent genomic markers to distinguish MSR from MSD may allow more accurate screening of the metabolic capacity of microorganisms from genome content alone in the absence of characterized isolates. Such markers have not yet been identified but are a target of active investigation (e.g., Umezawa et al., 2020). Finally, purely phylogenetic approaches to understanding the evolution of sulfur cycling in the Desulfobulbales provide an understanding of the timing of these processes only in relative evolutionary time. Tying this understanding to absolute, geologic time will require the application of additional approaches such as molecular clock analyses or calibrations using sediment geochemical and stable isotope records.

## Data Availability

All data utilized in this study is publicly available in the NCBI Genbank and WGS databases. Genomes first described here are available under submission ID SUB8971597 and will be released immediately following processing.

## Acknowledgements

Genomic DNA was acquired from the Deutsche Sammlung vonMikroorganismen und Zellkulturen (DSMZ). Genome sequencing was provided by MicrobesNG (http://www.microbesng.uk) which is supported by the BBSRC (grant number BB/L024209/1). EB and DTJ acknowledge support from NASA Exobiology (NNX15AP58G). This work was supported by a grant from the Simons Foundation (Grant Number 653687, LMW).

## Supplementary Material

### Tables

**Table S.1:**
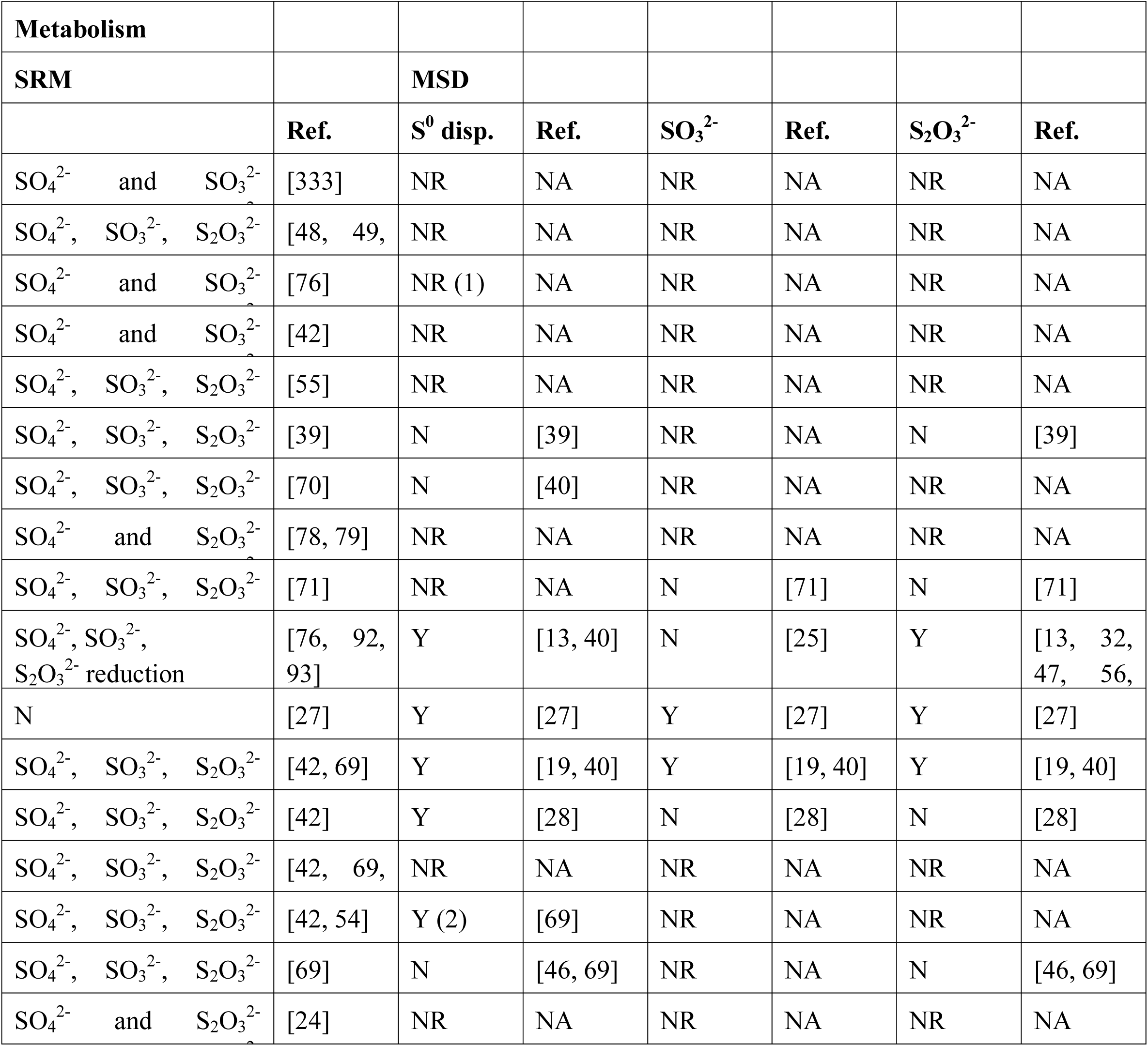
Summary of capacity to perform microbial sulfate reduction (type of electron acceptor used indicated) and/or microbial sulfur disproportionation (type of disproportionation performed indicated) as described in peer-reviewed reports. NR: Not Reported; NA: Not Applicable; (1) Only “personal communication” found then, by default, indicated as Not Reported; (2) Reported in the reference given but no supporting study found.

**Supplemental Table 2:**
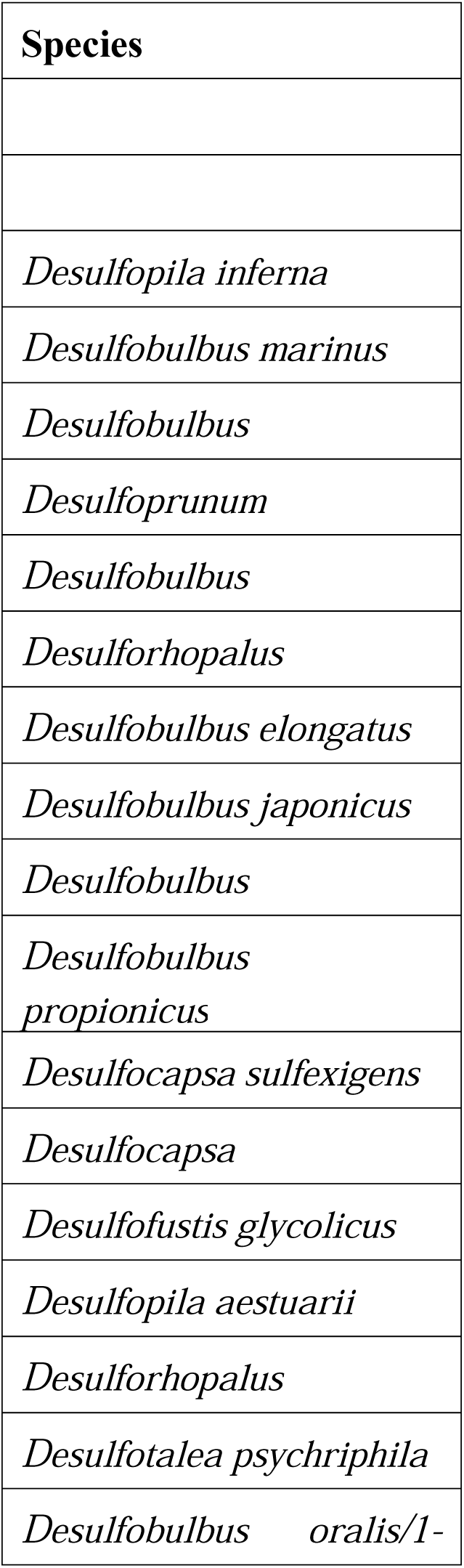
AAI matrix of Desulfobulbales

### Figures

**Figure S1:**
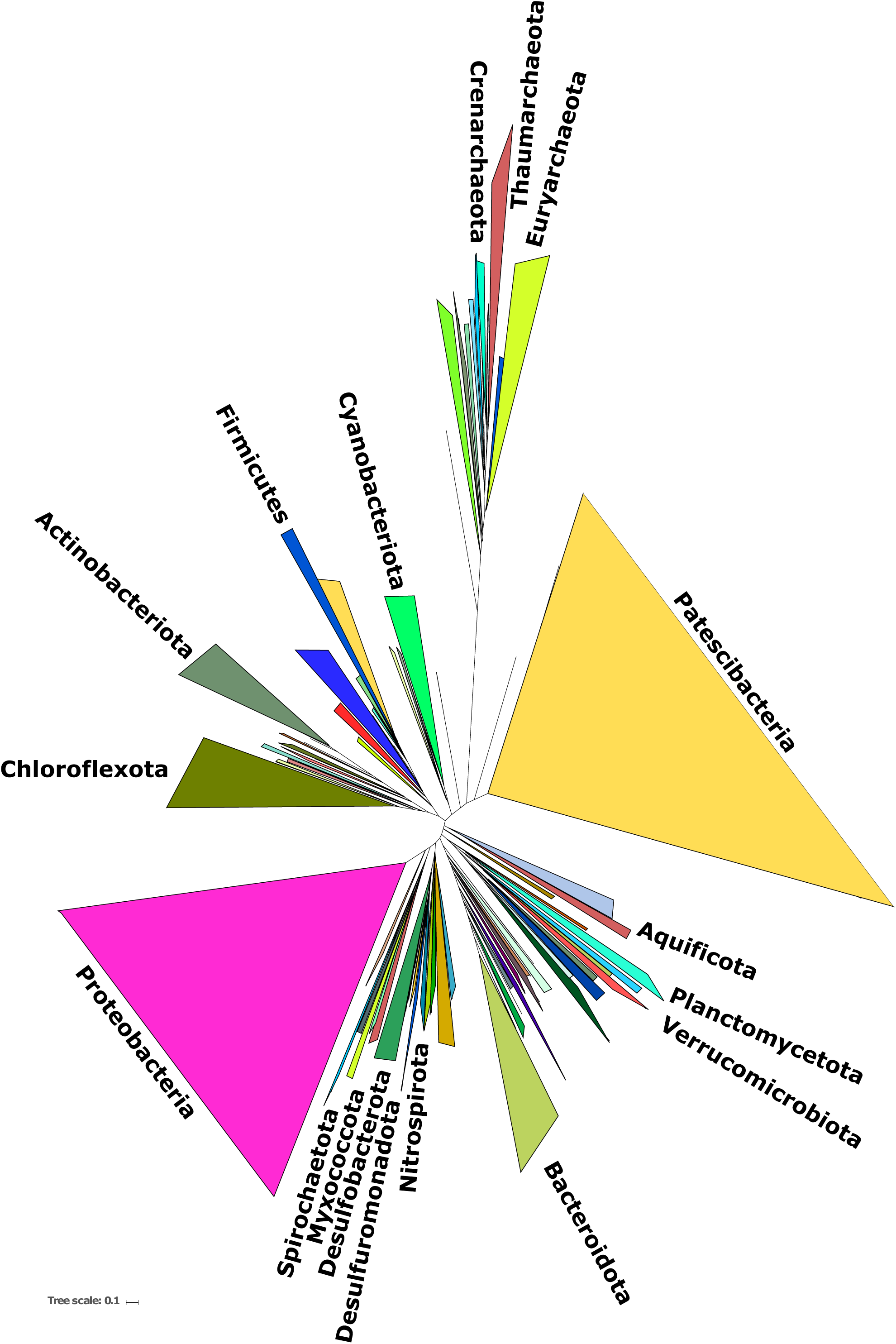
Phylogeny of Heme-Copper Oxidoreductase (HCO) proteins from members of the Desulfobacterota. Leaves are labeled with the family of HCO (A-, B-, or C-family) GTDB taxonomic assignments and WGS or Genbank IDs, nodes are labeled with TBE support value.

**Figure S2:**
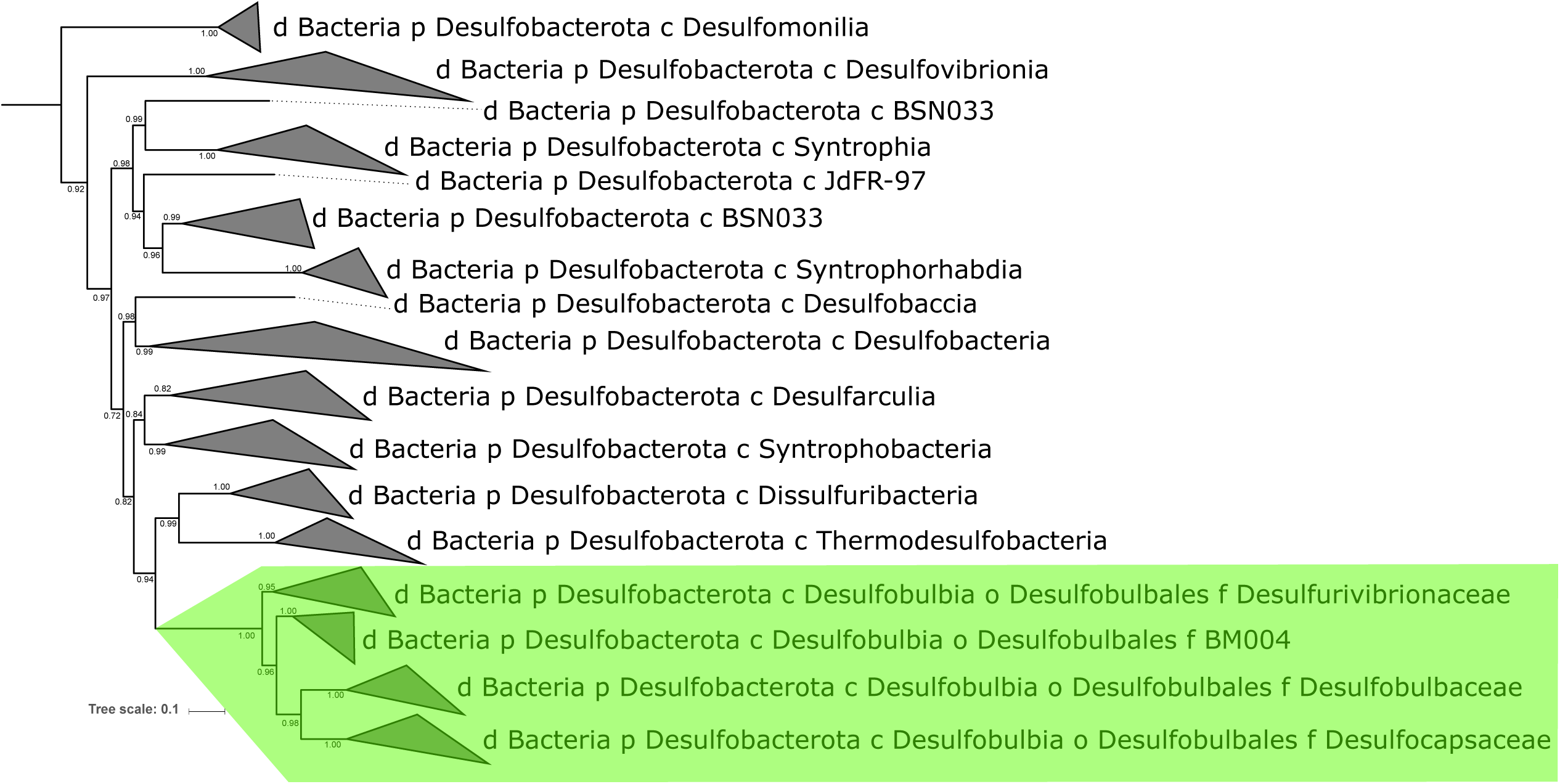
Phylogeny of concatenated DsrA, DsrB, DsrC, AprA, and AprB proteins from members of the Desulfobacterota. Leaves are labeled with GTDB taxonomic assignments and WGS or Genbank IDs, nodes are labeled with TBE support value.

**Figure S3:**
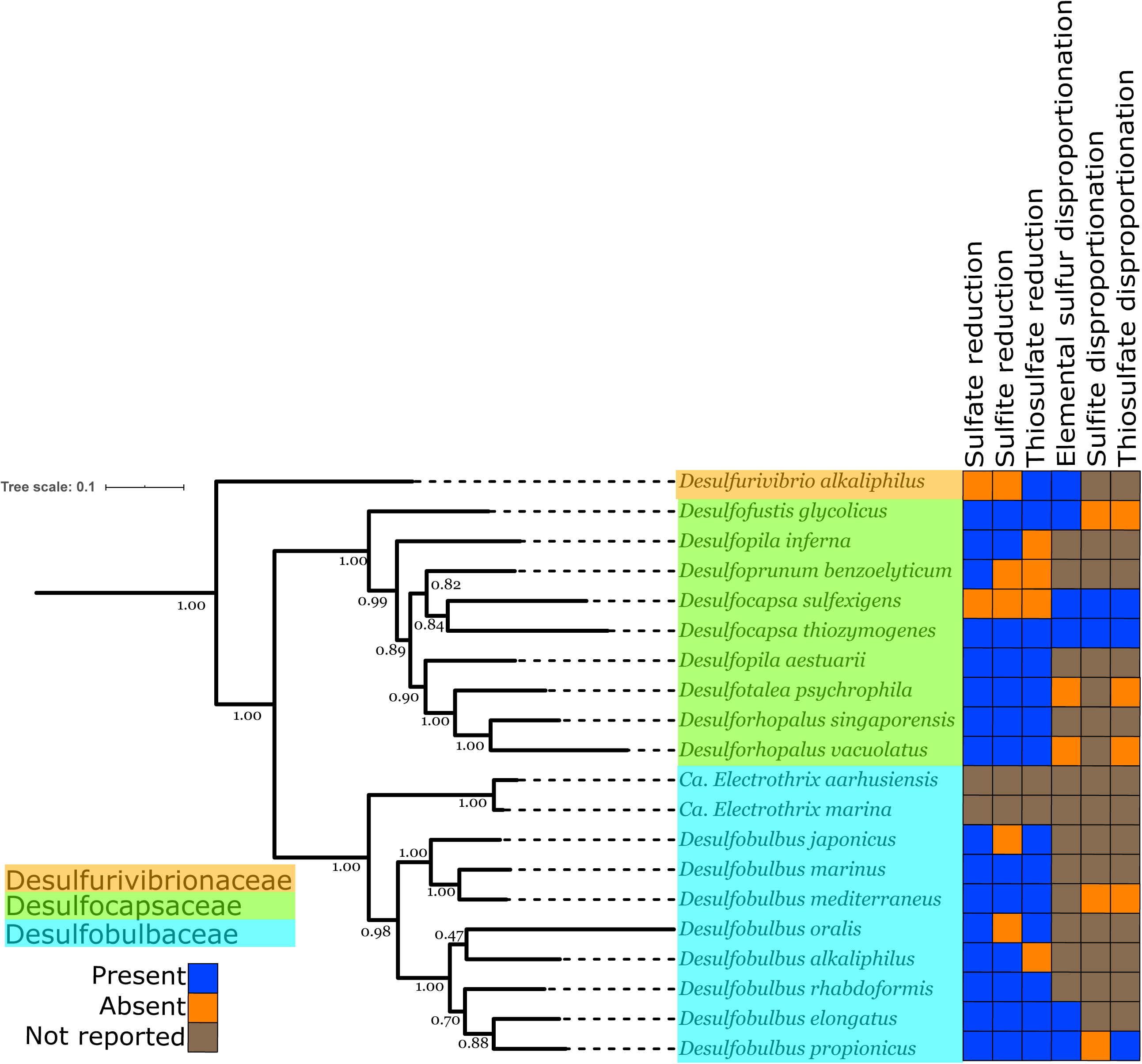
Phylogeny of *bd* oxidase proteins from members of the Desulfobacterota. Leaves are labeled with GTDB taxonomic assignments and WGS or Genbank IDs, nodes are labeled with TBE support value.

