## Supplementary figures and images for "Expanded Genomic Sampling of the Desulfobulbales Reveals Distribution and Evolution of Sulfur Metabolisms"

### Supplemental Figure 1

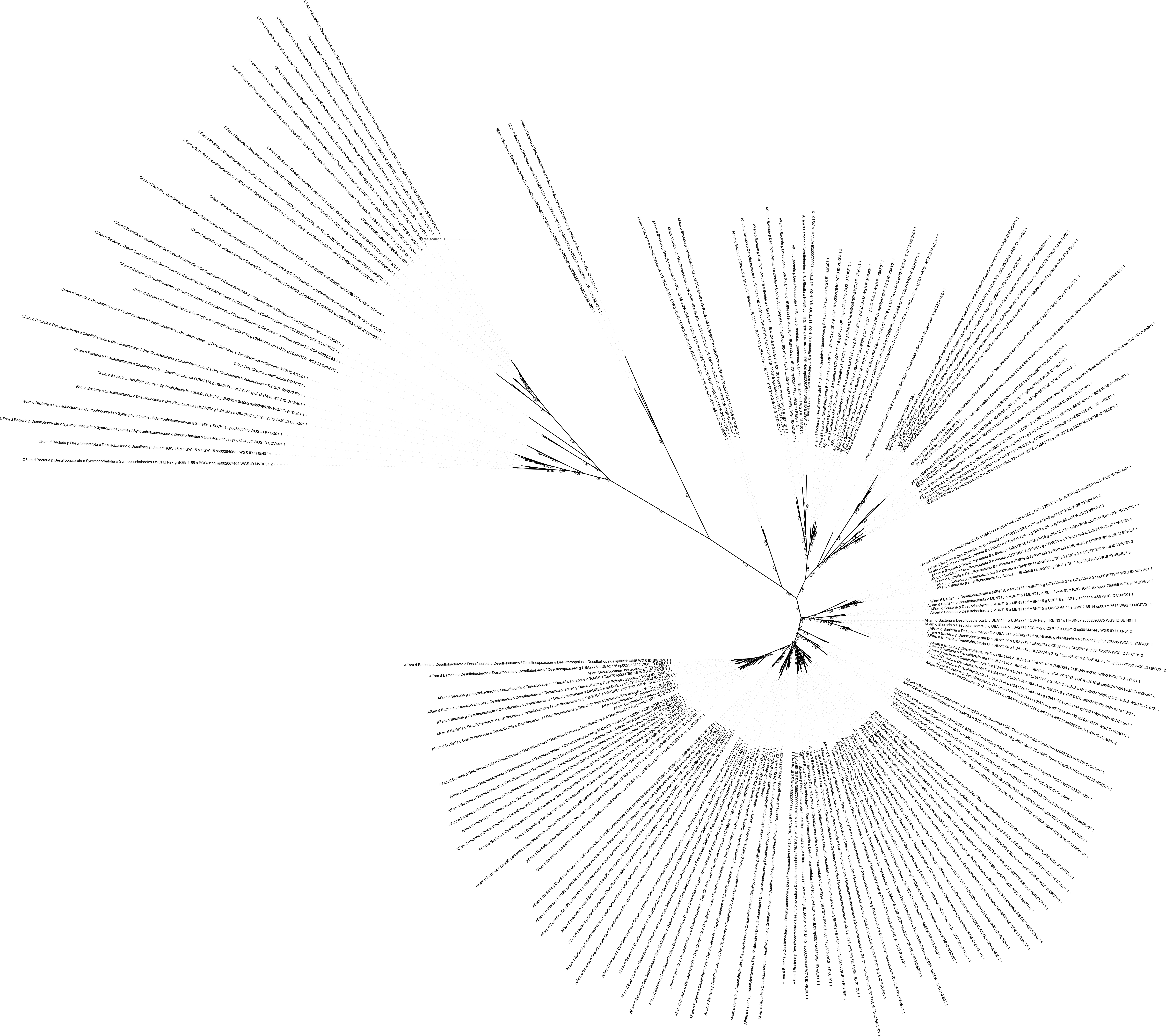

### Supplemental Figure 2

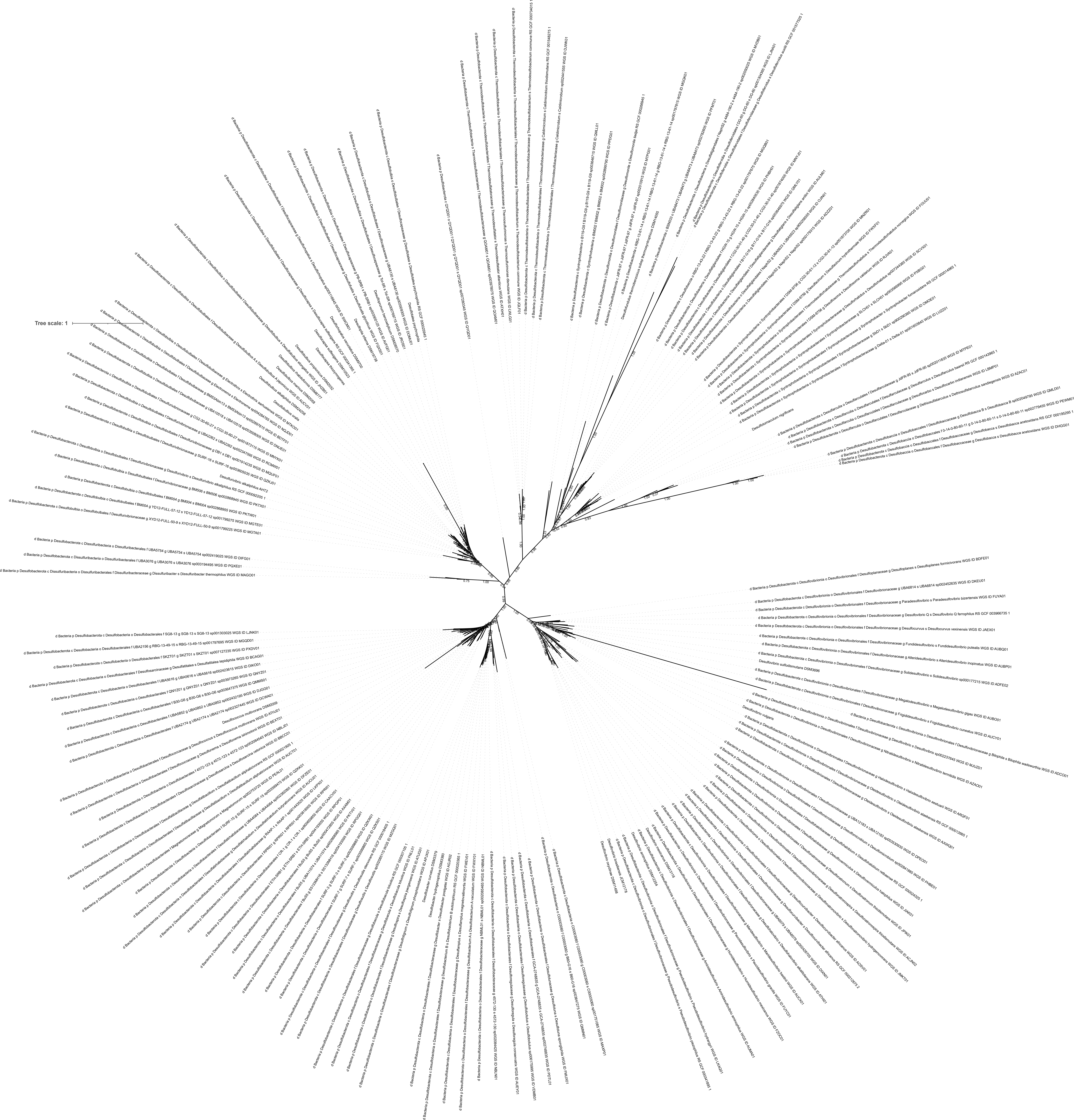

### Supplemental Figure 3

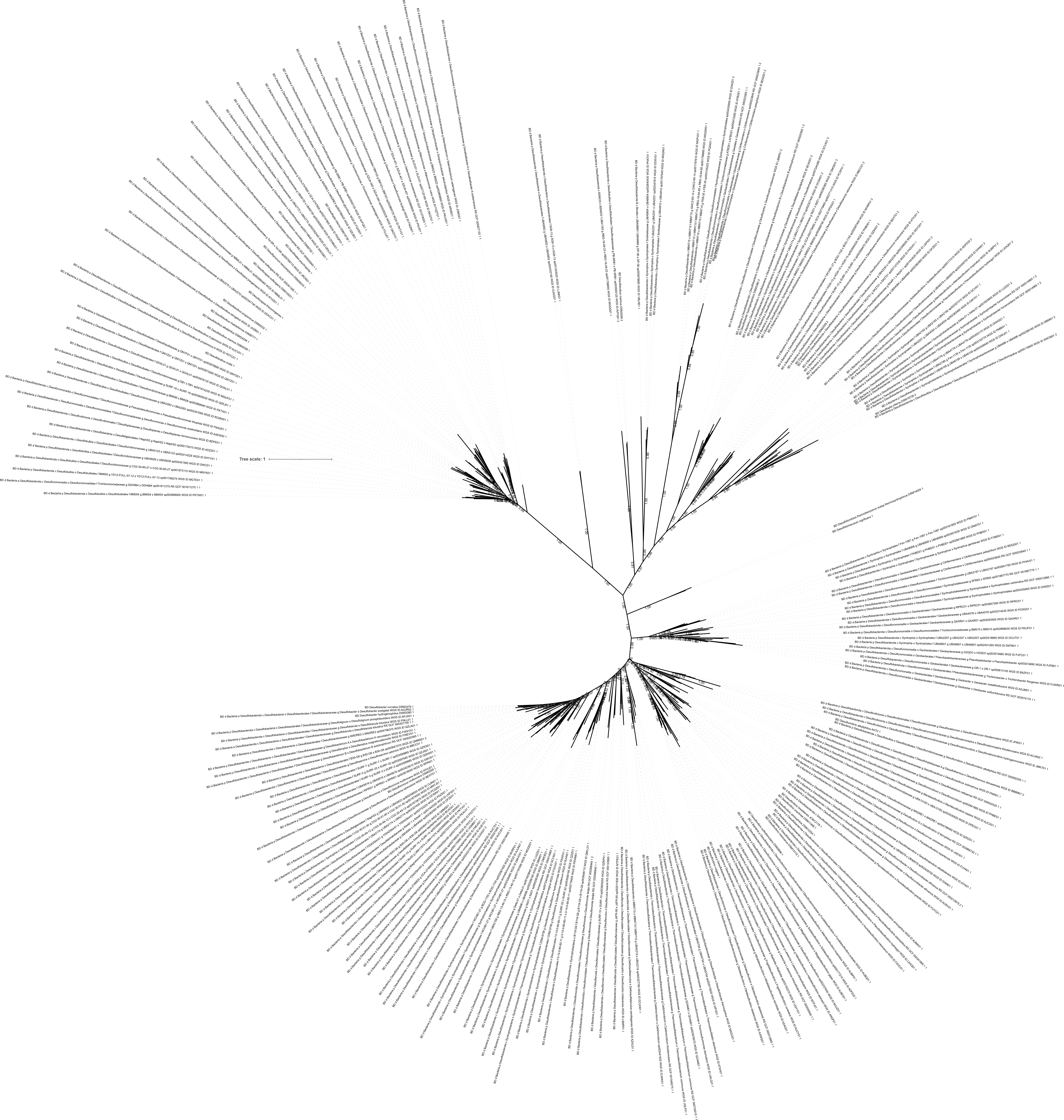
